# Green-synthesized silver and iron nanoparticles from *Alstonia boonei* and *Terminalia catappa as* antimalarial agents in mice models

**DOI:** 10.1101/2025.01.02.631173

**Authors:** Ojomona O. Abuh, Olabanji A. Surakat, Luqmon A. Azeez, Akinlabi M. Rufia, Kamilu A. Fasasi, Monsuru A. Adeleke

## Abstract

Malaria a major health scourge of high impact in sub-sahara, with a major control-drawback on issues of resistance of the parasite to antimalaria. Nanoparticles have been investigated for their capacity to effectively deliver antimalarial drugs to kill the parasites, avoid drug resistance evolution and maintain a low toxic side effects. This study investigated the antiplasmodial efficacies of two plant green-synthesized with AgNO_3_ and FeSO_4_ salts against *Plasmodium berghei* infected mice using, an acute toxicity test (LD_50_) of the *Terminalia-catappa* AgNPs (TCA), *Alstonia-boonei* AgNPs (AA), *Terminalia-catappa* FeSO_4_ (TCF) and *Alstonia-boonei* FeSO_4_ (AF) in accordance with Lorke’s method. The LD_50_ was above 5000 mg/kg of TCA and AA while 223.6 mg/kg in TCF and AF. Each nanoparticles exhibited high dose dependent *P. berghei* inhibition which was 62.20 % and 75.6 % in the TCA 200 mg/kg AA 100 mg/kg, 61.5 % and 75.3 % in the TCF 300 mg/kg and AF 200 mg/kg and doses in the curative groups, 92. 4 % and 93.1 % for TCA 100 mg/kg, AA 100 mg/kg 94.1 in 100 mg/kg dose and AF had 92.1 % at 300 mg/kg doses in the prophylactic groups. The groups treated with orthodox drugs had low parasitemia while the negative control recorded high parasitemia. There was no significant difference (P ˃ 0.05) in the mean change of the dosage in the different nanoparticles. These findings revealed all nanoparticles possesses a dose-dependent curative and prophylactic antiplasmodial activity and calls for their development and standardization as effective and readily available antimalarial options.

## Introduction

Malaria still constitutes a significant public health problem worldwide. The disease accounted for approximately 241 million malaria cases with over 620 000 deaths worldwide (WHO, 2021). Africa bore the brunt of the disease as over 95% of the cases and 96% of malarial deaths, with children under the age of five accounting for approximately 80% of malarial deaths occur in the region with profound impact on children under the age of five and pregnant women(WHO, 2021). This life threatening, parasitic disease is widely spread by the female Anopheles mosquito and its transmission are high during the wet season and could be all year round in many settings (WHO, 2021). Despite significant global efforts in the fight against malaria through increased funding for malaria research and development, delivery and scaling up of control interventions (diagnosis, prevention and treatment), the Global Technical Strategy (GTS) goals for malaria morbidity and mortality for 2020 are far from being achieved (WHO, 2018). Unfortunately, one of the major barriers to successful global malaria control (GMC) is the emergence and the propagation of parasites resistant to currently used antimalarial drugs. Artemisinin-based combination therapy (ACT), which is the most effective treatment available today, has been an integral part of the recent successes in GMC (White *et al*., 2014, WHO, 2018 and Ta *et al*., 2014). However, the future of these artemisinin based combinations is endangered by the emergence of artemisinin resistant *P. falciparum* (WHO, 2017, Souleymane *et al*., 2017 and Noedl *et al*., 2008). Effective malaria control and eradication depend largely on high-quality case management, vector control and surveillance (WHO 2006, 2012, 2014). Treatment with efficacious antimalarial drugs is crucial at all stages including the early control or ‘‘attack’’ phase to driving down transmission and the later stages of maintaining interruption of transmission, preventing reintroduction, and eliminating the last residual foci of infection (Bhatt *et al*. 2015; WHO 2007, 2014, 2016).

The future of malaria control and global elimination would depend on the ability of research and development to deliver the next generation of anti-malarial drugs (Tse *et al*., 2019). Diverse strategies exist for the development of novel anti-malarial drugs, and some have come from living organisms. Basically the synthesis of metal nanoparticles (NPs) requests the combination of three elements namely: the metal source (generally noble metals such as silver, gold, palladium and titanium salt), the reducing agent and the capping agent. Metal nanoparticles are traditionally produced using chemical and physical methods. However, these methods are challenging as they are costly, time-consuming and request for utilization of reagents harmful to environment (Eallas *et al*., 2017). In this regard, new NPs synthesis methods referred to as green synthesis have been developed to overcome these issues. Green synthesis consists in the production of metal NPs by exploiting the reducing and capping natural potential of biomolecules from living organisms such as plants and microorganisms. The method is simple, cost-effective and eco-friendly (Gahlawat *et al*., 2019). Nanoproducts and metal nanoparticles are highly useful, safe in nature with numerous applications in renewable energies, catalysis, cosmetics, food, electronics, environmental remediation, biomedical devices and health (Shakeel *et al*., 2016; Khandel *et al*., 2016). This study, investigated the efficacies of two plant (*Terminalia catappa* and *Alstonia boonei*) green-synthesized with AgNO_3_ salt for their antiplasmodial bioactive properties, efficacies and toxicity using mice model.

## Materials and methods

### Collections of Plant material

Fresh *T. catappa* and *A. boonei* leaves were collected in their full thriving stage around Omun, Coker Osogbo (Longitude: E 4.60388, Latitude: N 7.76117). They were validated in the taxonomy section of the Department of Plant Biology of Osun State University, Osogbo. The plants (plate 1a and b) were washed to remove dust, dried under room temperature for 7-14 days, the leaves were then collected and blended separately into fine particles and preserved in different airtight container till use. Other parts (Leaves, fruits, flowers and bark) of the plants were collected and taken to the Department of Plant Biology Laboratory, Osun state University, Osogbo for further identification, a voucher was deposited in the lab.

**Plate: 1.**
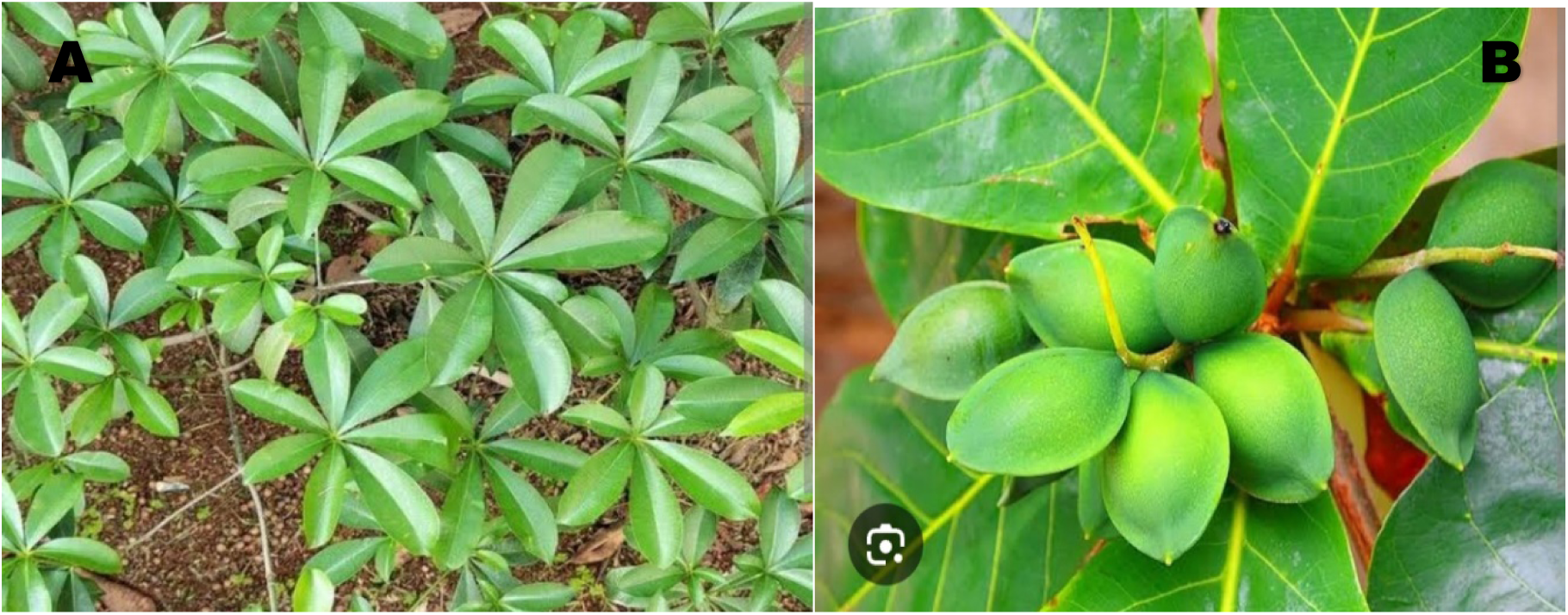
A. *Alstonia boonei* (God’s tree) B. *Terminalia catappa* (Almond)

### Preparation of plant extract

To prepare the aqueous extract, 10 grams of each of the already blended leaf extract was diluted with 1000 ml of distilled water, stirred allowed for 24 hours under room temperature, the solution was then be filtered using Whatman filter paper and refrigerated till use.

### Green-synthesis of silver nanoparticles

Green synthesis of AgNPs and FeNPs using *A. boonei* and *T. catappa* leaf extract was done following the procedures reported by Lateef *et al*., (2016). The reduction of silver ions was achieved by adding 7 mL of the extract with 293 ml of AgNO_3_ solution (1mM), while the reduction of ferrous ions was achieved by adding 10 mL of the extract with 1 ml of FeSO_4_ solution and both were incubated at room temperature The transparent brown color seen indicates the formation of silver nanoparticles while the amber colour in FeNPs gives the formation of iron nanaparticles. The solution was stored in a clean air-tight container and refrigerated until use.

### Characterization of Nanoparticles

#### UV-visible spectroscopy

The synthesized (TCA, AA, TCF and AF) nanoparticles was analyzed for Surface Plasmon Resonance with wavelength ranging from 200 - 1000 nm at room temperature using a UV-Visible spectrophotometer (Biobase BK-UV1900 P spectrometer, China)

### Fourier transform-infrared spectrometer (FTIR) analysis

The FTIR spectrum (TCA, AA, TCF and AF) synthesized nanoparticles was mediated to identify the characteristic functional groups of bioactive components using an infrared spectrum analyzer (SHIMADZU FTIR-8400S). The spectra were measured between 400 and 4000 cm1 in frequency.

### Scanning electron microscope (SEM) analysis

SEM (Phenom PRO X SEM-MVE01570775) was used to determine (TCA, AA, TCF and AF) synthesized nanoparticles size and shape. The constituent of the nanoparticles was analyzed with Energy Dispersive X-ray Fluorescence (EDXRF) Spectrum (ARL QUANT’X EDXRF Analyser. Serial No.9952120).

### Acute toxicity (LD_50_) test of *Terminalia catappa* AgNPs (TCA), *Alstonia boonei* AgNPs (AA), *Terminalia catappa* FeNPs (TCF) and *Alstonia boonei* FeNPs (AF)

A 50 % lethal dose (LD_50_) on the mice was performed on the two green synthesized nanoparticles treatments with *Terminalia catappa* AgNPs (TCA), *Alstonia boonei* AgNPs (AA), *Terminalia catappa* FeNPs (TCF) and *Alstonia boonei* FeNPs (AF) in accordance with Lorke (1983) with a slightly modification. This method involved the determination of LD_50_ value in biphasic manner. The animals were starved of feed but allowed access to water 24 h prior to the study. The tests were replicated in two phases, all the phases had six groups, and in each 1a, b and c and 2a, b and c which was treated orally, group 3a, b and c and 4a, b and c were treated intraperitoneally. A total of 144 mice were divided into two 72 mice in each, was used for this test. In the phase one, 72 mice divided into twenty four groups of 3 mice, were administered different doses (10 mg/kg, 100 mg/kg and 1000 mg/kg) of the test nanoparticles. The first three groups with TCA; the second three group AA; the third three groups TCF and the forth three groups AF respectively, the third and fourth three groups were intraperitoneally administered different doses (10 mg/kg, 100 mg/kg and 1000 mg/kg) of the test nanoparticles TCA, AA TCF and AF, and observed for 24 hours. In the phase 2 of the method, further specific doses were administered following same method as earlier stated (1600 mg/kg, 2900 mg/ kg and 5000 mg/kg) of TCA, AA, TCF and AF respectively. This was replicated for The LD_50_ was calculated as the square root of the product of the lowest lethal dose and highest non-lethal dose, i.e., the geometric mean of the consecutive doses for which 0 and 100% survival rates were recorded in the second phase, using the formula: LD50 = H (D0 9 D100); where D0 = highest dose that gave no mortality, D100 = lowest dose that produced 100% mortality (Lorke, 1983, Upkpanukpong *et al*., 2019 and Omagha *et al*., 2021).

### Antimalarial activities of *Terminalia catappa* AgNPs (TCA), *Alstonia boonei* AgNPs (AA) *Terminalia catappa* FeNPs (TCF) and *Alstonia boonei* FeNPs (AF)

The antimalarial potential of *Terminalia catappa* AgNPs (TCA), *Alstonia boonei* AgNPs (AA)*, Terminalia catappa* FeNPs (TCF) and *Alstonia boonei* FeNPs (AF) in mice infected with *Plasmodium berghei* was evaluated using both curative and prophylactic test models. The body weight of each mouse for all the tests was taken before and after exposure. 0.2 ml of the prepared *P. berghei* parasitized erythrocytes suspension in normal saline was injected intraperitoneally into each mouse to be used for the tests using 1 ml syringe and needle. The standard drugs (Chloroquine and Pyrimethamine), *Terminalia catappa* AgNPs (TCA), *Alstonia boonei* AgNPs (AA), *Terminalia catappa* FeNPs (TCF) and *Alstonia boonei* FeNPs (AF) were orally administered using an oral cannula. A daily parasitemia count was done using microscopic method of thin blood smears fixed in 100% methanol and stained with 10% Giemsa’s stain at pH 7.2 for 15 min were prepared from the tail blood of each mouse. Prepared blood films were allowed to air dry at room temperature and examined microscopically for malaria parasites under oil immersion (X100 magnification). The parasitaemia was determined by counting the number of parasitized erythrocytes in 2000 cells in randomly selected fields of the microscope. The percentage parasitaemia was determined as: Number of parasitized RBCs/Total number of RBCs (infected + Non-infected) × 100% (Fidock *et al*. 2004, Ukpanukpong *et al*., 2019). The mean parasite count for each group were determined and the percentage chemo inhibition for each dose was calculated as: [(A - B)/A], where A is the average percentage of parasitaemia in the negative control and B is the average percentage of parasitaemia in the test groups (Kalra *et al*. 2006).

### Rane curative test

The Rane curative test was conducted in accordance with Ryley and Peters (1970) in a similar method adopted by Omagha *et al*. (2021). Seventy-two hours after infection with chloroquine sensitive *P. berghei*, fifteen infected mice were divided into three groups of five each, each group were orally administered different doses (100, 200 and 300 mg/kg) of TCA, AA, TCF and AF respectively in the different 3 groups of mice. Fifteen mice were again divided into three groups of five each, the first five was infected but not treated as negative control, the second group was administered chloroquine phosphate (25 mg/kg) as positive control, while the last group received 10 mg/ml distilled water as the normal control. Each mouse was treated orally once daily with the dose given according to its body weight for four consecutive days (D4–D7) post inoculation during which the parasitaemia level were monitored daily.

### Repository (prophylactic) test

The method of Peters (1965) was adopted as reported by Omagha *et al*., (2019). Fifteen infected mice were divided into three groups of five each, each group were orally administered different doses (100, 200 and 3 00 mg/kg) of TCA. The same treatment was replicated for AA, TCF and AF respectively using another 3 groups of the animals. Fifteen mice were again divided into three groups of five each, the first five was infected but not treated as negative control, and the second group was given 5 mg/kg of pyrimethamine as the positive control, while the last group received 10 mg/ml distilled water as the normal control for four consecutive days. On the fourth day (D4), the mice were inoculated with *P. berghei*. Seventy two hours later (D7), smears were made from the mice (D7-D11) to check parasitaemia levels.

### Data analysis

Data collected from anti-plasmodial curative and prophylactic bioassays were entered and analyzed using Microsoft Excel version 2013 and Statistical Package for Social Sciences (SPSS) version 23.0. The differences between means among negative and positive controls as well as treatment groups were compared for significance using one way analysis of variance (ANOVA), followed by Duncan’s multiple post hoc test. Differences were considered significant to negative control at Probability ˂ 0.05.

## Results and Discussion

### Green-synthesis and characterization of AgNPs synthesized from both *A. boonei* and *T. catappa*

After an incubation period, a color change from yellow to transparent brown was seen, which implies bio-reduction of Ag^+^ to Ag^0^ (plate 2). The change in the appearance of the solution is a function of different bio-active constituents in the extract which serve as a reducing agents. According to Kanwal *et al*., (2019), the many stages of nucleation and development that occur during synthesis are visible in the color variation that occur overtime.

The spectrum of the biosynthesized AgNPs with an aqueous extract of *A. boonei* was monitored under UV-visible spectra at a wavelength between 190-700 nm. The absorption peak is characteristics to Surface Plasma Resonance (SPR) was recorded at 435 nm. The peak is indicative of reduced Ag and falls within the range characteristics of AgNPs (Figure 2). The functional groups of bioactive constituents in the biosynthesized AgNPs are shown in Figure 4. The prominent band observed were 3256, 3924, 2353, 1559, 1385 and 1072 cm^-1^. This bands were related to O-H stretching of alcohols, C-H alkane (CH stretching), C≡C, C=C stretching characteristics of alkenyl, CH_3_ bending absorption of methylene group and nitrogen group respectively (Figure 1 b-c). The biosynthesized nanoparticles were capped and synthesized by these functional groups. Fine whitish-ash cloudy pattern of AgNPs was observed in the SEM-EDs images, with the major components of the EDs patterns as Ag (77.20 %), O (10.20 %), N (8.30 %), Si (2.20 %) and S (2.10 %). The obvious presence of Ag in the EDs pattern implies the formation of AgNPs was confirmed (Figure 1d).

**Plate: 2.**
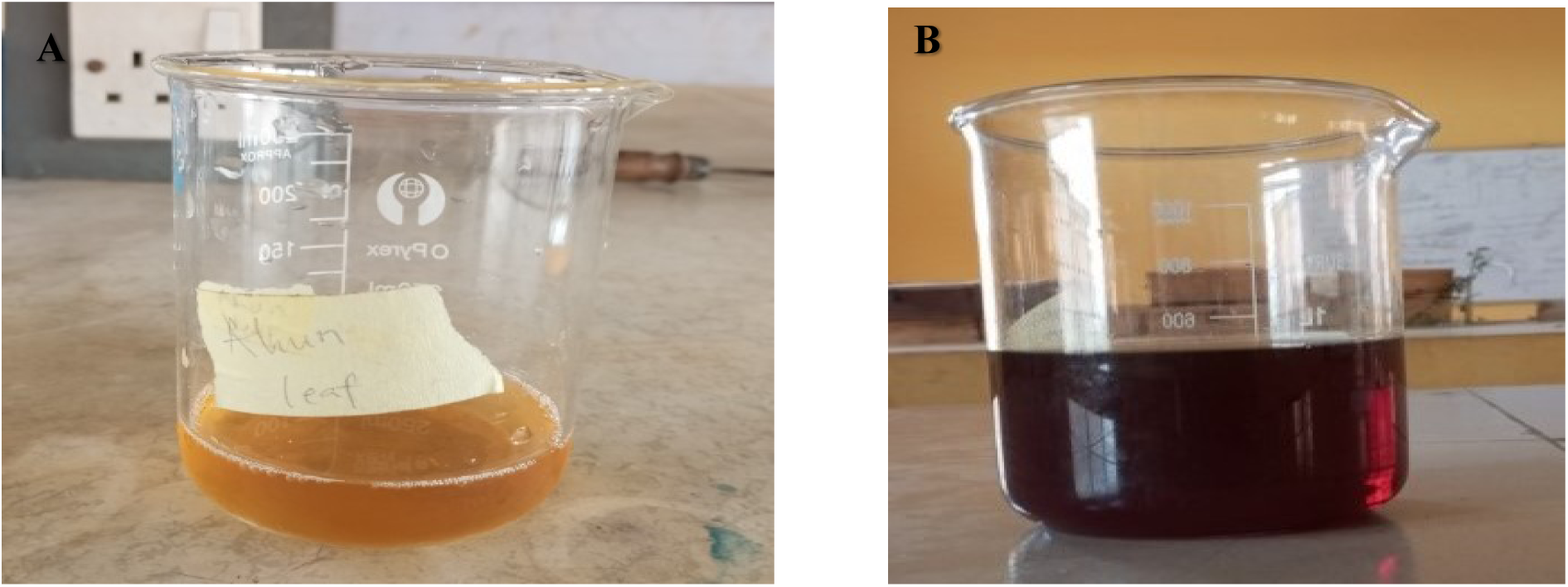
Color change of Biosynthesized AgNps of *A. boonei* leave extrac

**Figure: 1a-d.**
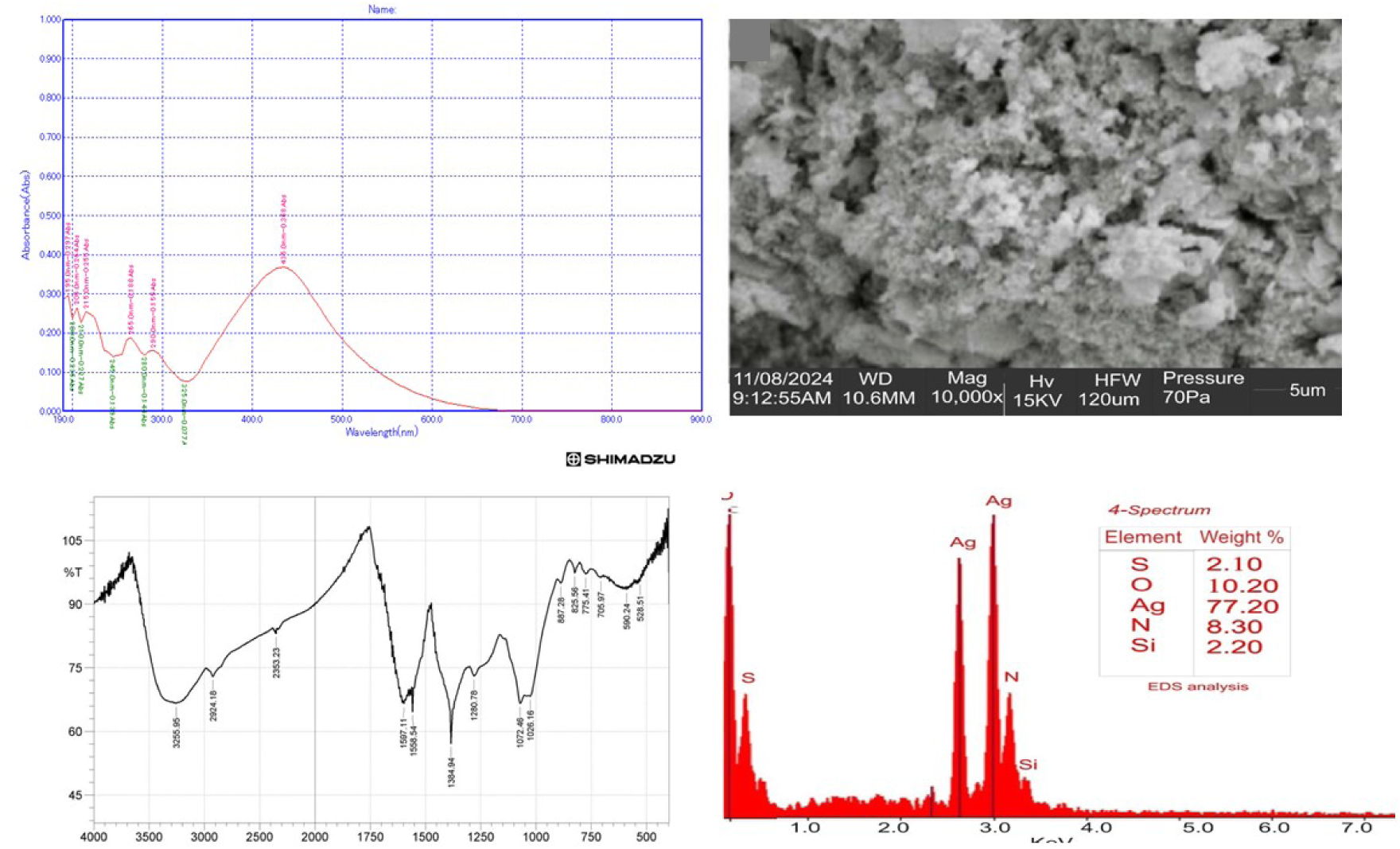
UV- spectrum; **b.** FTIR Spectroscopy; **c.** SEM image; **d.** EDS spectrum of AgNPs of *A. boonei*

### Green-synthesis of *T. catappa* with AgNO_3_

Upon incubation visual observation of gradual color change from light yellowish brown to dark brown was seen. This observation showed bio-reduction of Ag^+^ to Ag^0^. The change in the appearance of the solution is a function of the different bioactive constituent in the extract which serve as reducing agents (plate 3).

**Plate: 3.**
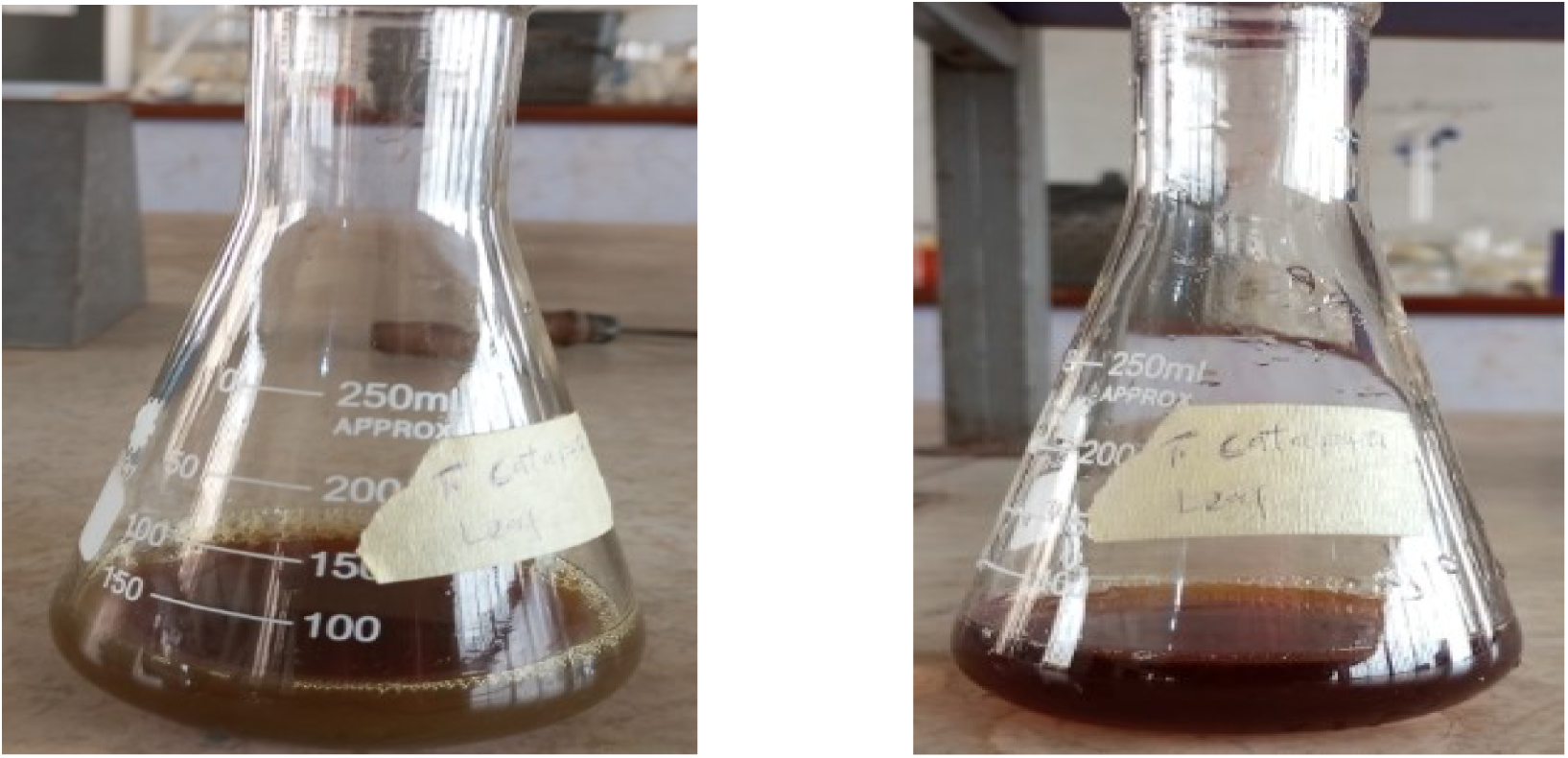
Color change of Biosynthesized AgNps of *A. boonei* leave extract

The spectrum of the biosynthesized AgNPs with an aqueous extract of *T. catappa* was monitored under the UV-Visible spectra at a wavelength between 190 - 700 nm (figure 2 a). The absorption peak characteristics of surface plasmon resonance (SPR) was recorded at 295 nm. This peak is indicative of reduced Ag and falls within the range characteristic of AgNPs (Ceylan *et al*., 2021: Azeez *et al* 2022: Aremu *et al*., 2023). It has been previously reported that an SPR peak between 410 and 450 nm as obtained in this study is related to spherical AgNPs.

The functional groups of bioactive constituents in a biosynthesized AgNPs are shown in (figure 8). The prominent bands observed were 3418, 3133, 2342, 1763 and 1395 cm^-1^. These bands were related O-H stretching and bending of primary and secondary alcohol, C-H stretching (alkane), C=O polyphenol and nitrate group respectively. The bio synthesized nanoparticles were stabilized and capped by these functional groups (Figure 2 b-c). The SEM-EDS images showed free irregular shape patterns of AgNPs, the main component of EDS patterns are Ag (65.20 %), O (20.20 %), N (9.20 %), Si (2.20 %), C (2.10 %), K (1.10 %). The overwhelming presences of Ag in the EDS pattern indicates the formation of AgNPs (Figures 2 d).

**Figure: 2a-d.**
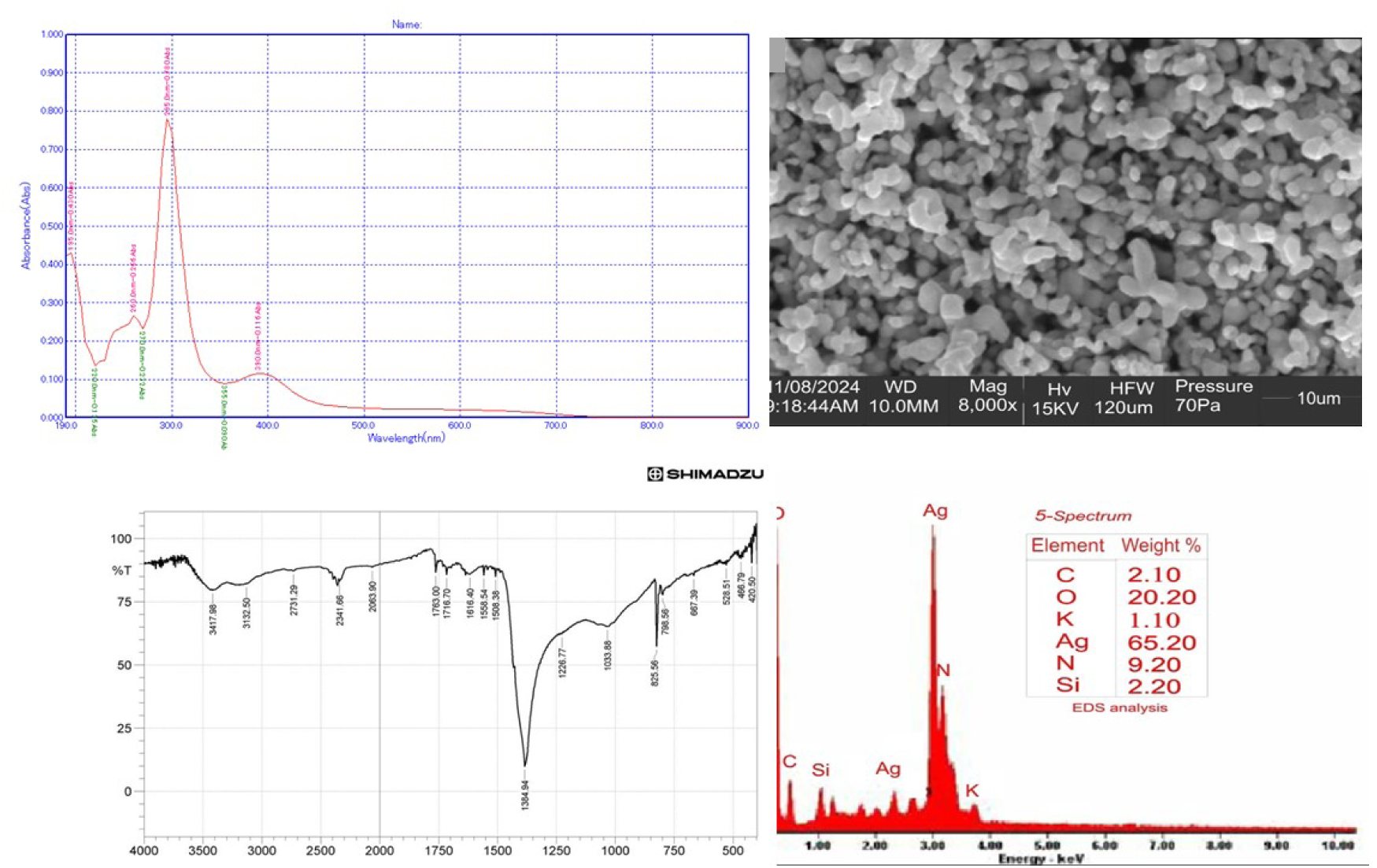
UV- spectrum; **b.** FTIR Spectroscopy; **c.** SEM image; **d.** EDS spectrum of AgNPs of *T. catappa*

### Green-synthesis and characterization of FeNPs synthesized from both *A. boonei* and *T. catappa*

#### Green-synthesis of *A. boonei*

After incubation over a period of time, the color change from light golden-yellow to deep amber transparent brown was observed (Figure 3a) which suggest bio-reduction of Fe^+^ to Fe^0^ the change in the appearance of the solution is a function of the different bioactive constituent in the extract which serve as reducing agents. The many stages of the nucleation and development that occur during synthesis are visible in the color variation that occur over time (Kanwal *et al*., 2019). The spectrum of the biosynthesized FeNPs with an aqueous extract of *A. boonei* was monitored under UV-visible spectra at a wavelength of between 190-900 nm (figure 3b) the absorption peak is characteristics to surface plasma resonance (SPR) was observed at 242 nm. This peak is indicative of reduced Fe and falls within the characteristics of FeNPs (Ceylan *et al*., 2021; Azeez *et al*., 2022). The functional groups of bioactive constituent in the synthesized FeNPs are shown in (Figure). The prominent bands observed were 3391, 2928, 1628 and 1080 cm^-1^. These bands were related to OH stretching of alcohol with C-H broad stretch of alkane, C=C and sulphate group, the presence of polyphenol protein as the functional group is responsible for capping and stabilizing (Alavi and Karimi 2018, Thakur *et al*., 2019 Azeez *et al*., 2021). The SEM-EDS image showed bunches of clouded pattern of FeNps. The major component of the EDS pattern are Fe (60.24 %), O (20.22 %), S (7.32 %), C (4.70 %), Ca (3.32 %), Na (2.20 %) and Si (2.00 %). The eminence of Fe in the EDS pattern indicates the formation of FeNps (Azeez *et al*., 2021) (figure 3 c-d).

**Plate: 4.**
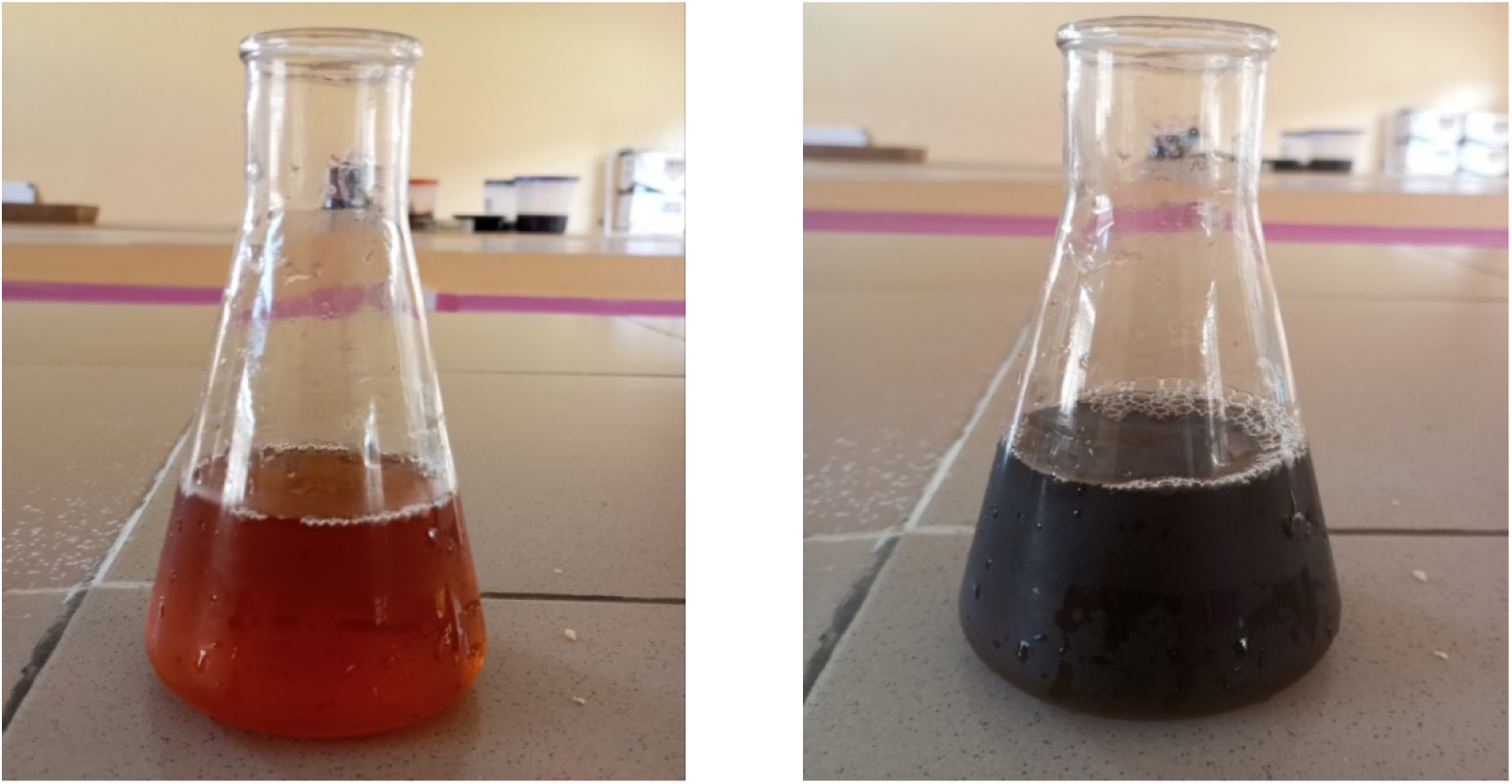
Color change of Biosynthesized FeNps of *A. boonei* leave extract

**Figure: 3a-d.**
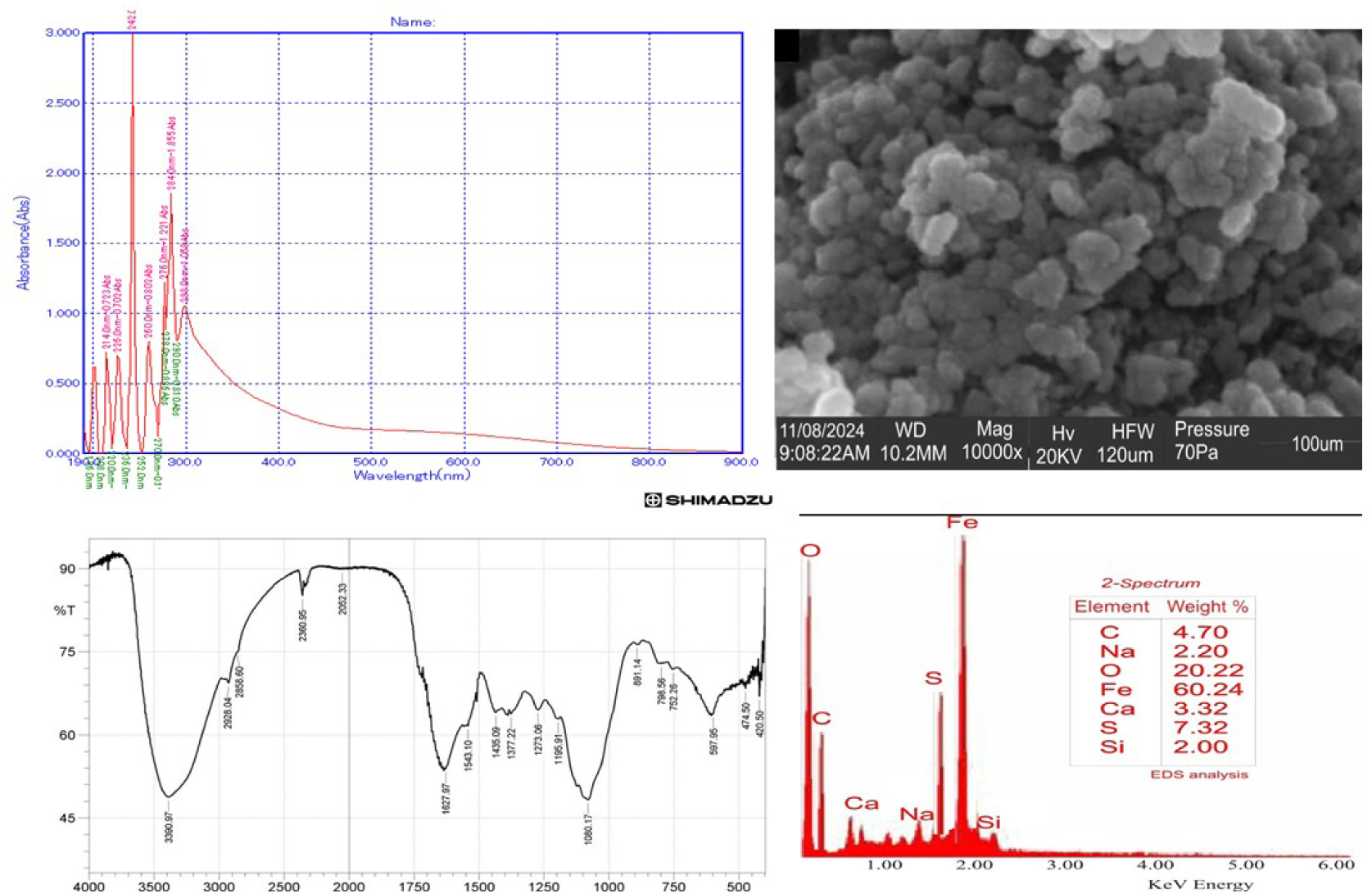
UV- spectrum; **b.** FTIR Spectroscopy; **c.** SEM image; **d.** EDS spectrum of FeNPs of *A. boonei*

**Figure: 4a-d.**
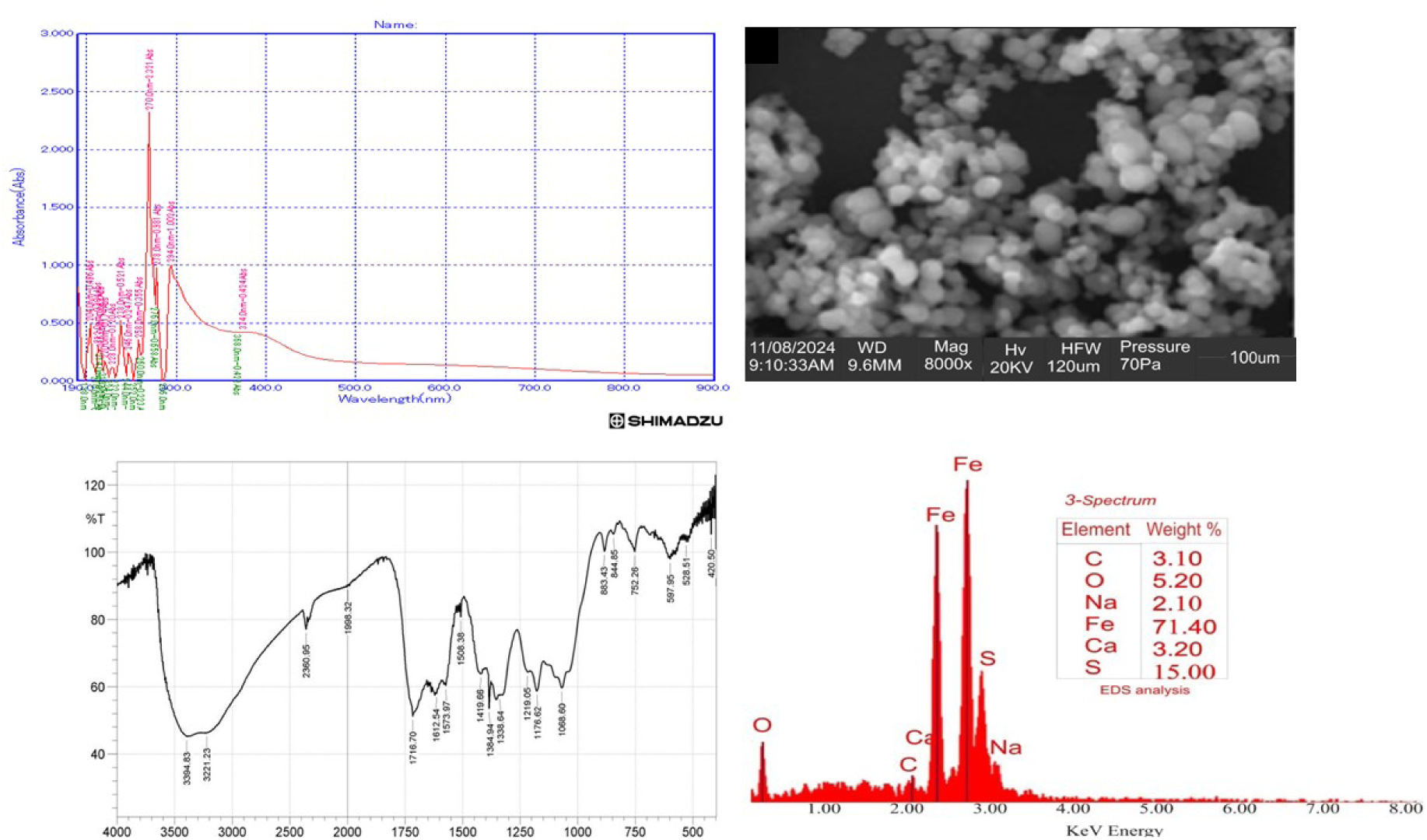
UV- spectrum; **b.** FTIR Spectroscopy; **c.** SEM image; **d.** EDS spectrum of FeNPs of *T. catappa*

#### Green-synthesis of *T. catappa*

After the incubation period, a color change from deep yellow/ burnt orange color to blackish-brown deep was seen which implies biosynthesis of Fe. The change in the appearance of the solution is a function of the different bioactive constituents in the extract which serve as reducing agents (Kanwal *et al*., 2019). The UV-visible spectra was observed at a wavelength of between 190-900 nm. The absorption peak is characteristic to surface resonance (SPR) was observed at 270 nm (figure 4a). This peak is indicative of reduced Fe and falls within the characteristic of FeNPs (Ceylan *et al*., 2021; Azeez *et al*., 2022). FTIR spectrum for *T. catappa* medicated FeNp_s_ manifested strong peaks at 3394.83 and 1716.70cm^-1^ implicating OH and C=O stretch of an ester or a carboxylic acid, The 1176.62cm^-1^ is a C-O stretching vibration while 1068.60 is a sulphate group respectively. The SEM-EDS image revealed a clustered cloudy pattern of FeNps. The main components of the EDS patterns are Fe (71.40 %), S (15.00 %), O (5.20 %), Ca (3.20 %), C (3.10 %) and Na (2.10 %) (figure 4 b-d). The strong presence of Fe in the EDS pattern relates to the presence and formation of FeNPs (Jyoti *et al*., 2016).

### Acute toxicity (LD_50_) test of *Terminalia catappa* AgNPs (TCA), *Alstonia boonei* AgNPs (AA) *Terminalia catappa* FeNPs (TCF) and *Alstonia boonei* FeNPs (AF) in mice model

The acute toxicity test of both groups administered with AgNPs-*Alstonia boonei*, AgNPs-*Terminalia catappa* FeNPs-*Terminalia catappa* (TCF) and FeNPs-*Alstonia boonei* (AF) revealed dose-dependent activity reduction, the mice were restless upon administration and slowly calm down and became sluggish after a few hours, mice in phase two were observed to be very sluggish and quite clogging together. After twenty four hours, no mortality was observed in the TCA, AA and TCF groups administered intraperitoneally and orally respectively, but one mortality was recorded in the group orally administered with 500 mg/kg with AF. Therefore the safe dose for TCA, AA and TCF is above 5000mg/kg while for AF is 223.6 mg/kg.

### In vivo antimalarial potencies of *Terminalia catappa* AgNPs (TCA), *Alstonia boonei* AgNPs (AA), *Terminalia catappa* FeNPs (TCF) and *Alstonia boonei* AgNPs (AF) in mice model

Upon the establishment of the parasites in the already infected mice, the curative effect were observed and calculated for four consecutive days, in all the treated groups there was a steady reduction in the parasitemia level from day 4 to day 7. Parasite inhibition was almost 100% with CQ 25 mg/kg from day 2 till day 5. In the treated groups, inhibition was observed to be 57.2 %, 62.2 % and 51.3 % in the TCA 100 mg/kg, TCA 200 mg/kg and TCA 300 mg/kg doses respectively; 75.6 %, 70.1 % and 60.6 % in the AA 100 mg/ kg, AA 200 mg/kg and AA 300 mg/kg doses respectively; 49.6 %, 58.7 % and 61.5 % in the TCF 100 mg/kg, TCF 200 mg/kg and TCF 300 mg/kg doses respectively; and 67.0 %, 75.3 % and 56.6 % in the AF 100 mg/ kg, AF 200 mg/kg and AF 300 mg/kg doses respectively.

Results of the Prophylactic effect on residual infection shows inhibition with PY 5 mg/kg to be 97.8 % throughout the 5 days of observation. With the nanoparticles treated groups, at the beginning of observation (D7), parasite inhibition was 93.1 %, 90.7 % and 92.1 % for the TCA 100 mg/kg, TCA 200 mg/kg and TCA 300 mg/kg doses respectively; 92.4 %, 91.7 % and 87.9% for AA 100 mg/ kg, AA 200 mg/kg and AA 300 mg/kg doses respectively; 94.1 %, 89.3 % and 89.0 % for the TCF 100 mg/kg, TCF 200 mg/kg and TCF 300 mg/kg doses respectively; and 91.7 %, 92.1 % and 92.4 % for AF 100 mg/ kg, AF 200 mg/kg and AF 300 mg/kg doses respectively.

**Figure 6a.**
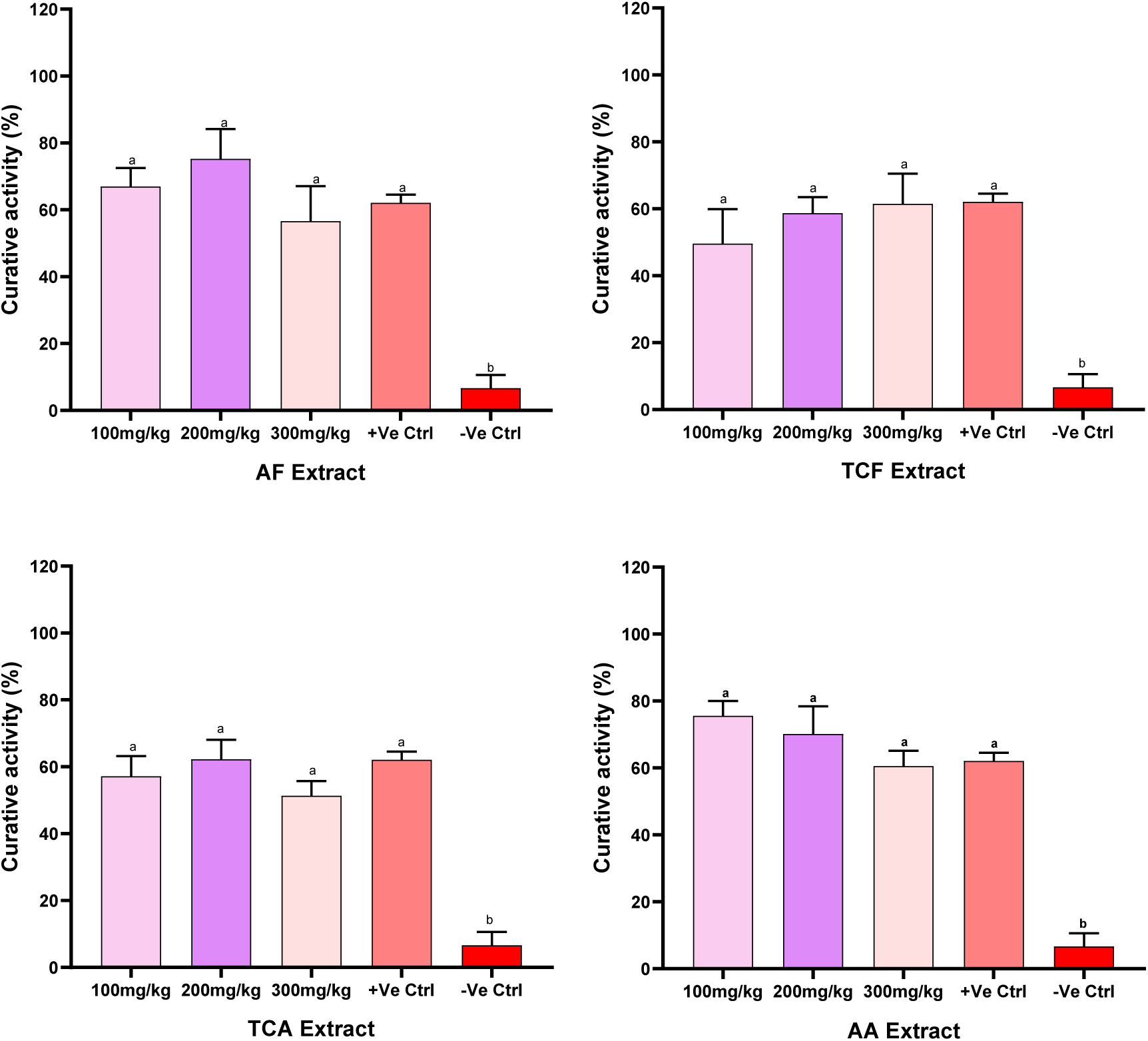
Curative activities of treatments with TCA, AA, TCF and AF on established infection

**Figure 6b.**
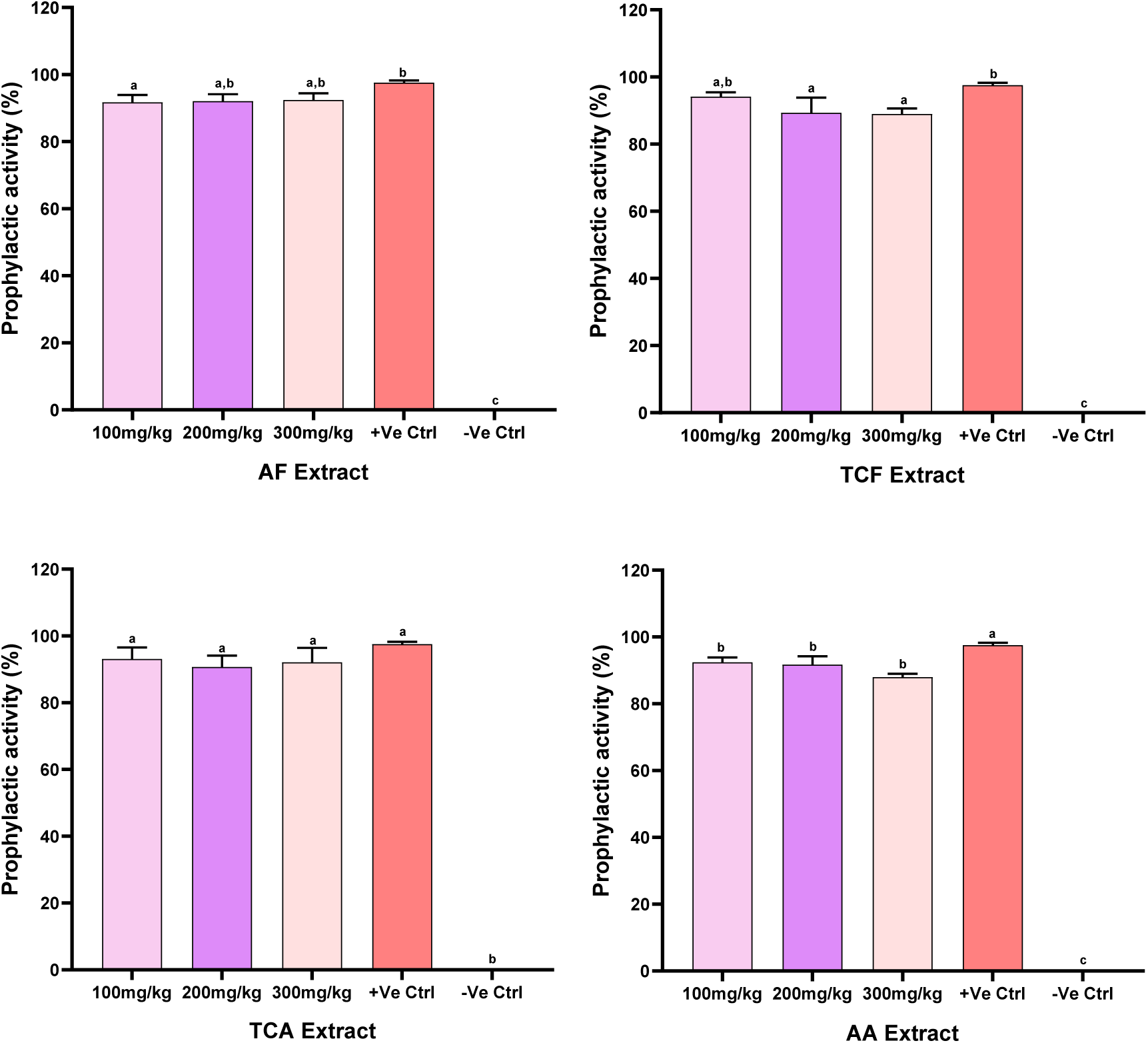
Prophylatic activities of treatments with TCA, AA, TCF and AF on residual infection

**Figure 6c.**
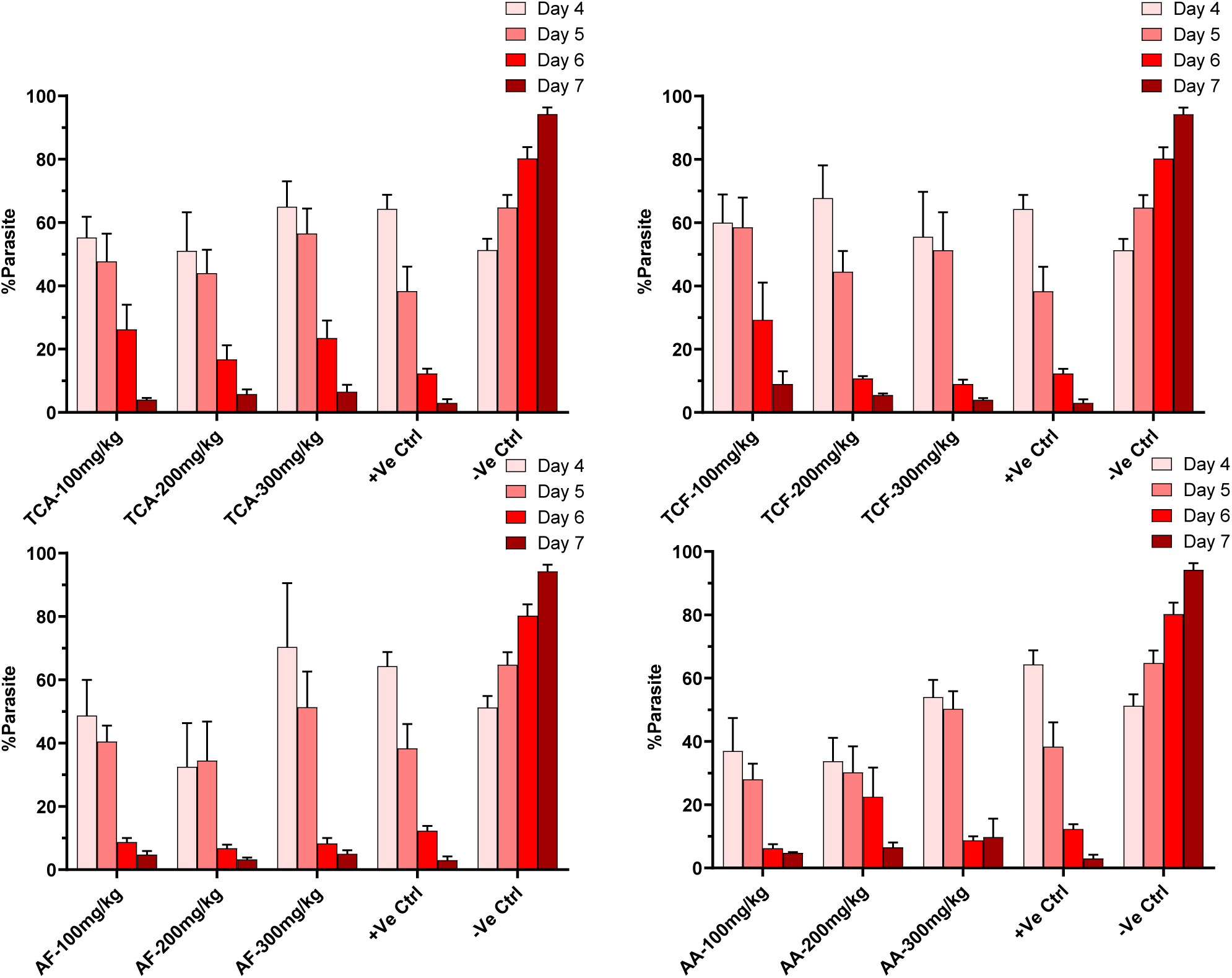
Curative activities of daily treatments with TCA, AA, TCF and AF on established infection

**Figure: 7a.**
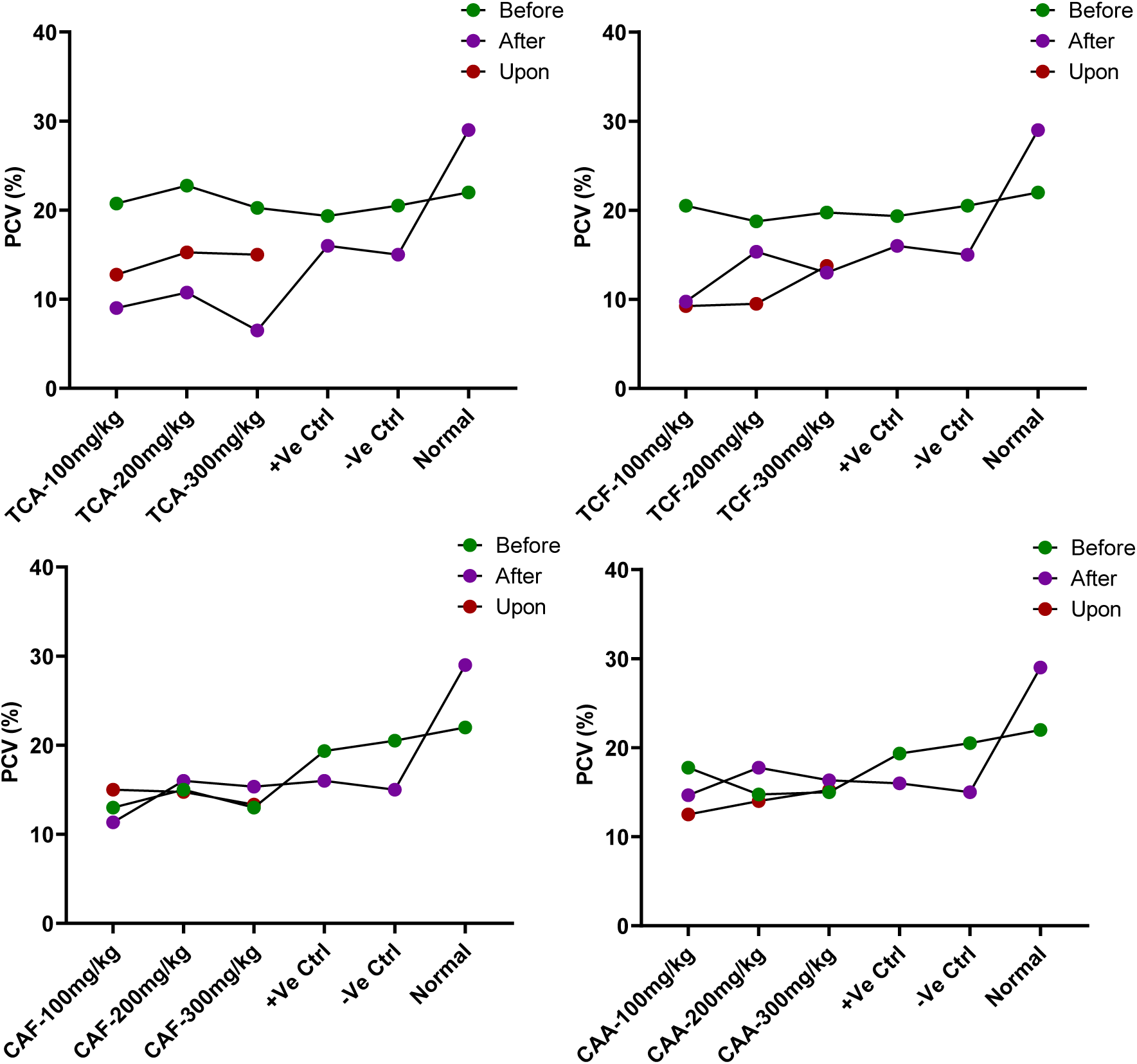
Packed cell volume of curative test model of before, upon and after inoculation and treatment with TCA, AA, TCF and AF

**Figure: 7b.**
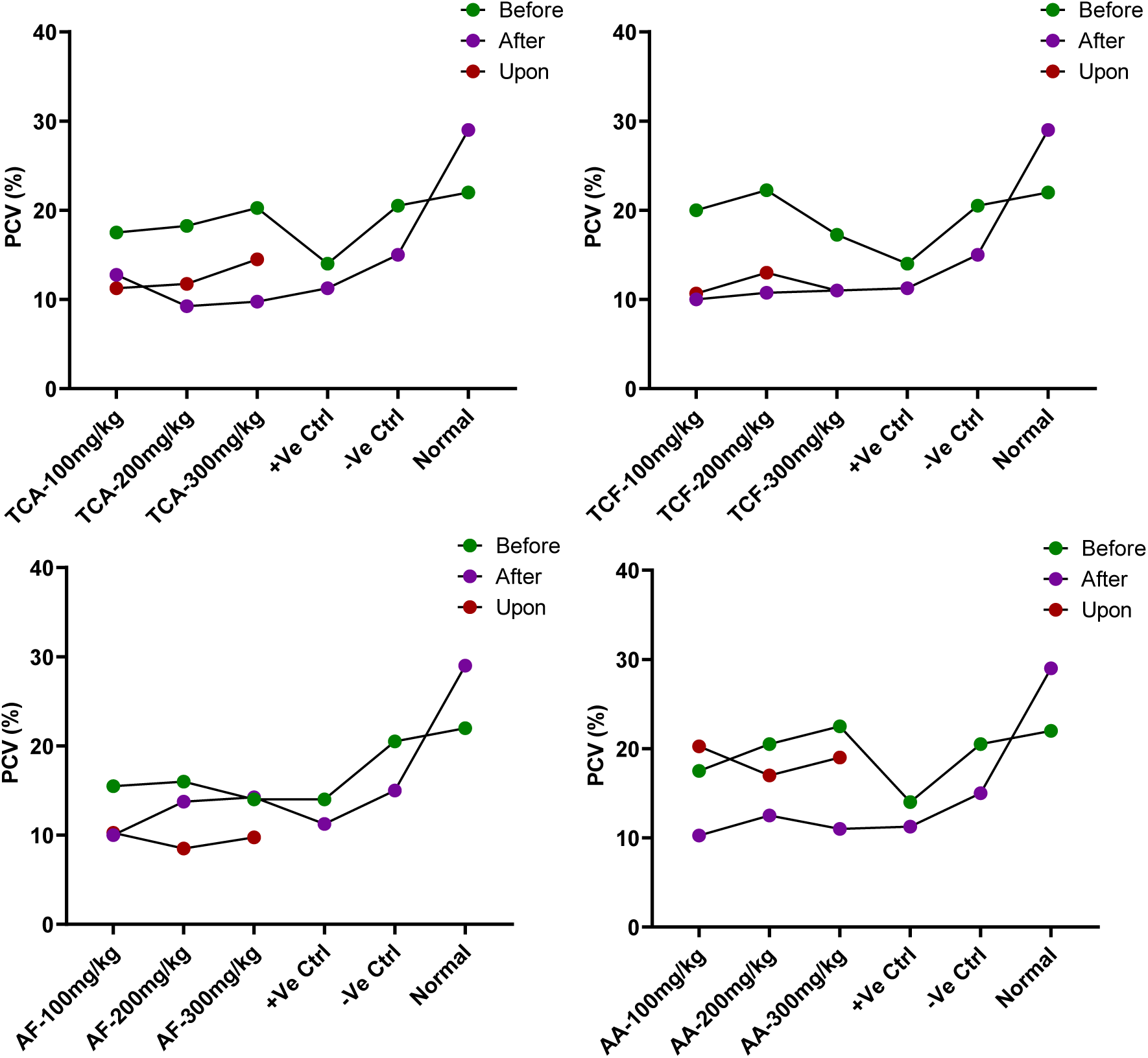
Packed cell volume of prophylactic test model of before, upon and after inoculation and treatment with TCA, AA, TCF and AF

## Discussion

The leaves extract of *A. boonei* and *T. catappa* was used to green-synthesize silver and iron nanoparticles (AgNPs and FeNPs) with a stable deep-brown and amber coloration, as reported in previous studies (Lateef *et al*., 2016; Azeez *et al*., 2017; Sabir *et al*., 2022; Aremu *et al*., 2023). The maximum absorbance of the fabricated AgNPs as analyzed under UV–visible spectra at wavelengths ranging from 190 - 700 nm had a strong peak at 435.00 nm, which agrees with Yusuf-Omoloye *et al*., (2024) a value suggestive of the reduction in Ag, and clearly within the range of several previously reported qualities of spherical silver nanoparticles (AgNPs) of SPR values between 410 to 450 nm (Chinnathambi *et al*., 2023; Lateef *et al*., 2016; Aremu *et al*., 2023, Busi *et al*., 2014; Saheed *et al*., 2020; Azeez *et al*., 2022). While the maximum absorbance of the fabricated FeNPs as analyzed under UV–visible spectra at wavelengths ranging from 190 - 900 nm with the absorption peak is characteristics to surface plasma resonance (SPR) was observed at 270 nm (TCF) and 242 nm (AF), a value suggestive of the reduction in Fe. Furthermore, FTIR analyses revealed the vibrational kinetics of atoms or molecules, as well as absorption peaks. The prominent band observed were 3256, 3924, 2353, 1559, 1385 and 1072 cm^1^. This bands were related to O-H stretching of alcohols, C-H alkane (CH stretching), C≡C, C=C stretching characteristics of alkenyl, CH_3_ bending absorption of methylene group and nitrogen group respectively. (Alabi and karimi 2018. Thakur *et al*., 2019, Azeez *et al* 2021, Aremu *et al*, 2023). Also the FTIR spectrum for *T. catappa* medicated FeNp_s_ manifested strong peaks at 3394.83 and 1716.70cm^-1^ implicating OH and C=O stretch of an ester or a carboxylic acid, The 1176.62cm^-1^ is a C-O stretching vibration while 1068.60 is a sulphate group respectively (Alabi and karimi 2018. Thakur *et al*., 2019, Azeez *et al* 2021, Aremu *et al*, 2023). While, The functional groups of bioactive constituent in the synthesized *Alstonia boonei*-FeNPs revealed prominent bands at 3391, 2928, 1628 and 1080 cm^-1^. These bands were related to OH stretching of alcohol with C-H broad stretch of alkane, C=C and sulphate group, the presence of polyphenol protein as the functional group is responsible for capping and stabilizing (Alavi and Karimi 2018, Thakur *et al*., 2019 Azeez *et al*., 2021). The prominent bands observed were 3418, 3133, 2342, 1763 and 1395 cm^-1^. These bands were related O-H stretching and bending of primary and secondary alcohol, C-H stretching (alkane), C=O polyphenol and nitrate group respectively (Alavi and Karimi 2018; Thakur *et al*., 2019, Azeez *et al*., 2021; Aremu *et al*., 2023). The biosynthesized nanoparticles were stabilized and capped by these functional groups. The functional groups of bioactive components in the fabricated AgNPs and FeNPs by *A. boonei* and *T. catappa* leaves aqueous extract, which are primarily responsible for the reduction, stabilization, and capping of the established nanoparticles. Although different plant extracts have been used to synthesize AgNPs and FeNPs, specific patterns of the peaks for each previously reported biosynthesized AgNPs (Aremu *et al*., 2023; Alavi *et al*., 2018; Dash *et al*., 2020) fall in sync with our observed peak pattern. The AgNPs had a clustered crystalline pattern, and the EDS pattern revealed Ag as the most abundant element (77.20 and 65.20 % respectively), indicating that the nanoparticles formed were mostly silver (Ag). An EDS peak of an additional metal, C, O, K, N and Si, was observed, but it was very low, which could be attributed to plant element composition. It can be deduced that the quality of nanoparticles produced is dependent on the characteristic and richness of phytochemicals in the extract used (Chinnathambi *et al*., 2023) as the phytochemical component in plants is the major component reducing the silver nitrates (AgNO_3_) into AgNPs (Barabadi *et al*., 2020; Lateef *et al*., 2016). For the SEM-EDS image revealed a clustered cloudy pattern of FeNPs. The main components of the EDS patterns are Fe (71.40 %), S (15.00 %), O (5.20 %), Ca (3.20 %), C (3.10 %) and Na (2.10 %). The strong presence of Fe in the EDS pattern relates to the presence and formation of FeNPs (Jyoti *et al*., 2016) While the SEM-EDS *Alstonia boonei*-FeNPs image showed bunches of clouded pattern of FeNps. The major component of the EDS pattern are Fe (60.24 %), O (20.22 %), S (7.32 %), C (4.70 %), Ca (3.32 %), Na (2.20 %) and Si (2.00 %). The eminence of Fe in the EDS pattern indicates the formation of FeNps (Azeez *et al*., 2021) indicating that the nanoparticles formed in the both were mostly Iron (Fe). An EDS peak of an additional metal, S, O, Ca, C, Si and Na, was observed, but it was very low, which could be attributed to plant element composition.

This acute toxicity test was done to observe the mice for signs of toxicity including sluggishness, clogging together, weakness, anorexia, micturition, excitement, dullness, ataxia and mortality associated with administration of TCA and AA in mice model. The oral and intraperitoneal routes’ toxicity effects in this study showed no mortality within the different dosage concentrations. Thus, the LD_50_ of TCA, AA and TCF are above 5000 mg/kg respectively, while, the lethal dose for AF is 223.6 mg/kg. All the toxicity signs showed are negligible or insignificant toxicity signs (Okokon *et al*., 2017; Zemicheal and Mekonnen, 2018). Though intraperitoneal administration is sometimes preferred over the oral route as high concentration of the drug is able to bypass the physiological barriers and provide the highest bioavailability and the fastest effect (Cardenas *et al*. 2017), administering large volumes intraperitioneally (10 ml/kg in rodents) can lead to pain, chemical peritonitis, formation of fibrous tissue, perforation of abdominal organs, hemorrhage, and respiratory distress (Bredberg *et al*., 1994; Esquis *et al*., 2006). These concerns of potential adverse effects may necessitate more favorable routes of administration. Therefore, these nanoparticles are not toxic. This result is similar to that obtained by Nkono, for the study of the acute toxicity of the aqueous extract of the stem bark of *A. boonei* collected at Ombessa in Cameroon (Nkono *et al*., 2014). The results are also similar to those obtained by Iyiola *et al*. (2011) and Dibua *et al*. (2013) on the *A. boonei* leaves ethanolic extract collected in Shagari and Nsukka (Nigeria).

The two models (curative and prophylactic), results revealed that TCA, AA, TCF and AF significantly inhibited parasitaemia, with inhibition levels above to 50 % in TCA above 60 % in AA, above 49 % in TCF and 56 % in AF of the 100, 200 and 300 mg/kg doses of each nanoparticles in the curative models, while over 90 % inhibition level in the prophylactic model in all the different doses. AA at 100 mg/kg had the highest inhibition, TCA at 200 mg/kg showed the highest inhibition, TCF at 300 mg/kg had it highest inhibition and AF at 200 mg/kg had its highest inhibition for the curative model, while for the prophylactic model for both AA, TCA and TCF 100 mg/kg had the highest mean parasitemia inhibition while AF at 300 mg/kg had its highest mean parasitemia inhibition when compared to all the other treated groups. The daily parasitemia level showed a steady reduction at doses 100 mg/kg in TCA and AA while 300 mg/kg in TCF and 200 mg/kg in AF respectively as compared to the other treated groups respectively. The results of the antiplasmodial for curative and prophylactic activity showed a dose-dependent reduction in mean parasitemia of the mice, the least parasitemia was observed in the group treated with the orthodox drugs whereas the negative control group had the highest parasitemia. Hence, the mean parasitemia in relation to the treatments showed a very high significant difference. These findings confirms antiplasmodial potency of TCA, AA, TCF and AF providing scientific experimental basis for their usage as antimalarial remedies and proves they are potential sources that should be explored for new antimalarial drugs. Chloroquine, an antimalarial drug still widely used in Nigeria because it is accessible and affordable, despite WHO recommendation for Artemisinin Combination Therapies ACTs (WHO 2001a; b; Oladipo *et al*. 2015) was used as the reference drug in the curative models (Ukpanukpong *et al*., 2019; UIwalokun 2008; Alli *et al*. 2011), and Pyrimethamine was the standard antimalarial drug used in the prophylactic model (Alli *et al*., 2011 Omagha *et al*., 2021). These standard drugs inhibited parasitaemia to an almost undetectable levels and the findings agrees with results from other studies validating medicinal plants in malaria treatments (Alli *et al*. 2011; Ogbole *et al*. 2014). The antiplasmodial activity observed in these nanoparticles could be attributed to the presence of some of the antimalarial proven phytochemicals including saponin, alkaloid, flavonoid, phenol and lactones (Kirby *et al*., 1989; Philipson and Wright 1991; Christensen and Kharazmi 2001; Onifade and Maganda 2015; Ibukunoluwa 2017; Omagha *et al*. 2020) present in each of the plants used in the green-synthesis of TCA, AA, TCF and AF. These phyto-constituents affect the condition and function of body organs, clear up residual symptoms of the disease. They help increase the body’s resistance to disease or facilitate the adaptation of the organism to certain conditions (Njan, 2012). The antiparasitic activity of the green-synthesized nanoparticles is largely dependent on the prompt production of phytochemicals and the response to radical mopping by the nanoparticles (Tighe-Neira *et al*., 2020; Borgheti-Cardoso *et al*., 2020) Nanomaterials, which have been extensively focused on cancer-related biomedical applications for many years, are being intensively investigated to implement strategies to prevent and cure malaria, with the aim of overcoming the problems arising with conventional therapies (Santos-Magalhães *et al*., 2010; Urban *et al*., 2014; Dennis *et al*., 2015). Nanocarriers are suited to be functionalized with several antigens, which might dramatically boost the prospects of obtaining a stronger immune response than with the current classical vaccination approaches (Borgheti-Cardoso *et al*., 2020) The decrease in PCV after inoculation agrees with the clinical manifestation of malarial parasites in rodents, the gradual increase in the PCV upon the treatment of all the infected groups after infections are in conformity with previous reports (Saganuwan *et al*., 2011; Abuh, 2022). In this study, administration of TCA and AA nanoparticles triggered an increase in PCV as a result of increased production of RBC, thus, suppressing hemolytic damage to RBC.

## Conclusion

Findings in this study demonstrates that TCA, AA, TCF and AF possess good antimalarial abilities and safer in both oral and intraperitioneal administrations. The results clearly indicate that the individual administration of TCA, AA TCF and AF significantly decreased parasite load in mice, increased their PCV and enhanced their survival as compared to the negative control groups.

## Acknowledgements

thanks to the authority of the Department of Animal and Environmental Biology, University of Osun state University Osogbo, Osun state, Nigeria for providing some of the facilities and equipment used in the studies that have been cited in this article The authors are grateful to the Institute of Advanced Medical Research and Training (IMRAT), University College Hospital, Ibadan, Nigeria, for kindly supplying the rodent parasite, *Plasmodium berghei berghei* NK 65. E. Idowu and O. Otubanjo Department of Zoology, University of Lagos, E. Evans (Federal university of Technology Minna) O. Oyedara (Department of Biotechnology), E. Ajayi and H Aremu. of the Department of Biochemistry, are appreciated for their support. The Department of Pure and Applied Chemistry of Osun state University, Federal university of Technology Akure, Redeemers University Ede are also acknowledged for technical assistance.

## Authors’ contributions

Abuh O. O. and Adeleke M. A. conceptualized the idea for this study. Abuh O. O. wrote the research proposal. Adeleke M. A. Surakat O. A., Azeez L., reviewed the research proposal and contributed to improving the experimental design. Abuh O. O. conducted the research and drafted the manuscript. All the authors participated in reviewing and approving the work for publication.

## Declarations

Conflict of interest the authors declare that they have no conflict of interest.

## Ethical clearance

the ethical clearance was obtained from the Department of Health Planning, Research and Statistics, Ministry of Health, Osogbo, Osun State.

## Informed consent

This study involving laboratory mice for antimalarial experiments were carried out following Nigerian Institute of Medical Research Institutional Review Board’s guidelines for laboratory animal care and use.

